# Anti-GPR56 monoclonal antibody potentiates GPR56-mediated Src-Fak signaling to modulate cell adhesion

**DOI:** 10.1101/2020.05.11.089177

**Authors:** Treena Chatterjee, Sheng Zhang, Tressie A. Posey, Ling Wu, Wangsheng Yu, Qingyun J. Liu, Kendra S. Carmon

## Abstract

GPR56, also known as ADGRG1, is a member of the adhesion G protein-coupled receptor (aGPCR) family shown to play important roles in cell adhesion, brain development, immune system function, and tumorigenesis. GPR56 is upregulated in colorectal cancer and high expression correlates with poor prognosis. Several studies have shown that GPR56 couples to the Gα_12/13_ class of heterotrimeric G-proteins to promote RhoA activation. However, due to its structural complexity and lack of a high affinity receptor-specific ligand, the complete GPR56 signaling mechanism still remains largely unknown. To delineate the activation mechanism and intracellular signaling functions of GPR56, we generated a monoclonal antibody (mAb) which binds GPR56 with high affinity and specificity to the extracellular domain (ECD). We show that overexpression of GPR56 in 293T cells lead to increased phosphorylation of Src, Fak, and paxillin adhesion proteins and activation of the Gα_12/13_-RhoA-mediated serum response factor (SRF) pathway. Treatment of cells with anti-GPR56 mAb potentiated Src-Fak phosphorylation, RhoA-SRF signaling, and cell adhesion. Consistently, knockdown of GPR56 in colorectal cancer cells decreased Src-Fak pathway phosphorylation and cell adhesion. Interestingly, GPR56-mediated activation of Src-Fak phosphorylation occurred independent of RhoA, yet mAb-induced potentiation of RhoA-SRF signaling was Src-dependent. Furthermore, we show that the Serine–Threonine–Proline-rich (STP) region of the ECD of GPR56 was not essential for mAb binding, yet was required for activation of Src-Fak signaling. These data support a new ECD-dependent mechanism by which GPR56 functions to regulate cell adhesion through activation of Src-Fak signaling.

## Introduction

GPR56, also referred to as ADGRG1, is a member of the adhesion G protein-coupled receptor (aGPCR) family with a large N-terminal extracellular domain (ECD) that is thought to function in mediating cell-cell and cell-matrix interactions (1). GPR56 has been shown to be critical for many physiological processes, including immune function (2), maintenance of hematopoietic stem cells (3), and oligodendrocyte and cortex development (4–7). Mutations in GPR56 have been associated with bilateral frontoparietal polymicogyria (BFPP), a disorder characterized by a disruption in the organization of the frontal cortex (6,8). Recently, GPR56 has been implicated in contributing to tumorigenesis of various types of cancer. GPR56 is upregulated in cancers of the breast, lung, ovary, pancreas, colon and glioblastoma (9–11). Analysis of clinical data revealed a significant correlation between high GPR56 levels and poor outcome in acute myeloid leukemia, ovarian cancer, and colorectal cancer (CRC) (10–15). In CRC, GPR56 was shown to promote drug resistance and drive tumor growth (12,13,16). On the contrary, GPR56 has been shown to be downregulated in metastatic melanoma and inhibitory to melanoma growth and metastasis (17). These studies demonstrate the essential functions of GPR56 and its emerging, yet diverse roles in tumor progression.

Similar to other aGPCRs, the ECD of GPR56 is structurally characterized by the presence of a highly conserved <u>G</u>PCR-<u>A</u>utoproteolysis <u>IN</u>ducing (GAIN) domain featuring a juxtamembrane <u>G</u>PCR <u>P</u>roteolysis <u>S</u>ite (GPS) (1,18). The GPS can be autoproteolytically cleaved, leaving two non-covalently associated but distinct fragments: (1) the N-terminal fragment (NTF) which consists of a <u>P</u>entraxin and <u>L</u>aminin/neurexin/sex-hormone(LNS)-binding-globulin-<u>L</u>ike (PLL) domain with adhesion properties, an overlapping <u>S</u>erine-<u>T</u>hreonine–<u>P</u>roline-rich (STP) region, and the bulk of the GAIN domain; (2) and the C-terminal fragment (CTF) which incorporates the C-terminal region of the GAIN domain, referred to as the stalk or *Stachel*, and the seven-transmembrane region (7TM) (1,19). Typically, aGPCR activation involves binding of the NTF with its ligand followed by activation of the CTF to transmit intracellular signaling (1). Evidences supporting both stalk-dependent and stalk-independent models of ligand-induced GPR56 activation have been reported, however the exact mechanism remains elusive. In the stalk-dependent model, the NTF serves as a shield for the stalk region and upon ligand binding the NTF dissociates or sheds from the CTF, allowing stalk to function as a “tethered agonist” that can engage with the 7TM (20–22). Alternatively, the stalk-independent model suggests that a ligand-induced conformational change of the ECD (incorporating the NTF and the stalk region), which promotes interaction with the 7TM, is required to stimulate signaling activity (20,23). GPR56 has been shown to interact with proteins such as collagen III (24), transglutaminase 2 (TG2) (17), and progastrin (16); however receptor-specific ligands have yet to be fully validated. Several studies have established that GPR56 is coupled to the Gα_12/13_ class of heterotrimeric G-proteins to promote RhoA activation (13,22,24–26). GPR56 overexpression has been shown to activate various signaling pathway response elements, including serum response element (SRE) and serum response factor response element (SRF-RE) that are likely regulated downstream of RhoA (20,21,25). However, due to its structural complexity and lack of high affinity receptor-specific ligands, the activation and intracellular signaling mechanism(s) of GPR56 have yet to be completely resolved.

In this study, we generated a high affinity anti-GPR56 monoclonal antibody (mAb) 10C7 to investigate the activation and signaling mechanisms of GPR56. We show that GPR56 can activate Src-Fak adhesion signaling in 293T and CRC cells. GPR56-mediated activation of this pathway required the STP region of the ECD, indicating the importance of the NTF in activation of Src-Fak signaling. Treatment with 10C7 potentiated both RhoA-SRF and Src-Fak signaling. GPR56-mediated activation of Src-Fak occurred independent of RhoA. However, activation of Src was required for 10C7-induced potentiation of RhoA-SRF signaling. Overall, we demonstrate a new ECD-dependent mechanism by which GPR56 functions to regulate cell adhesion through activation of the Src-Fak pathway.

## Results

### Characterization of monoclonal antibody 10C7 targeted to the ECD of GPR56

We generated and purified a unique mouse-human chimera anti-GPR56 mAb 10C7 directed against the extracellular domain (ECD) of GPR56 (amino acids 1-400). To characterize 10C7, we first established HEK293T (293T) cell lines with stable overexpression of either myc-tagged full-length human GPR56 wild-type (293T-GPR56WT), mutant GPR56 lacking the STP region (a.a.108-177) of the ECD (293T-GPR56ΔSTP), or vector control (293T-vector). GPR56 expression in these cells lines was verified by western blot (Fig. 1A). The STP region encompasses the C-terminal portion of the PLL domain (a.a. 108-160), the PLL-GAIN linker (a.a.161-175), and first two amino acids of the GAIN domain (a.a. 176-177) (19,27). Previously, it has been shown that the STP region is required for GPR56 interaction with TG2 (27) and collagen III has been shown to interact within the PLL domain (a.a. 27–160) (24), suggesting this region plays an important role in ligand/protein interactions. To demonstrate 10C7 binding specificity and test if its epitope maps to the STP region, we performed immunocytochemistry (ICC). All three cell lines were treated for 1 hour at 37°C with 10C7 (5 μg/ml). ICC and confocal analysis showed 10C7 binds and promotes internalization of GPR56WT and GPR56ΔSTP in a similar fashion (Fig. 1B), suggesting that its epitope is not located within the STP region of the ECD. ICC using a Cy3-labeled anti-myc-tag mAb which binds the N-terminus of GPR56 showed a similar pattern of internalization (Fig. S1B). To determine relative binding affinities, we employed a whole cell-based fluorescence binding assay. We showed that 10C7 had equal and high affinity binding for both WT and ΔSTP with approximate K_d_ of 1.2 μg/ml (or 7.8 nM) and 1.1 μg/ml (or 7.5 nM), respectively (Fig. 1C). No binding of 10C7 was detected in vector cells or for human IgG1 isotype control (Fig. 1B-C and Fig. S1A). These findings indicate that 10C7 binds GPR56 with high affinity and specificity, yet does not bind within the STP region of the ECD.

**Figure 1.**
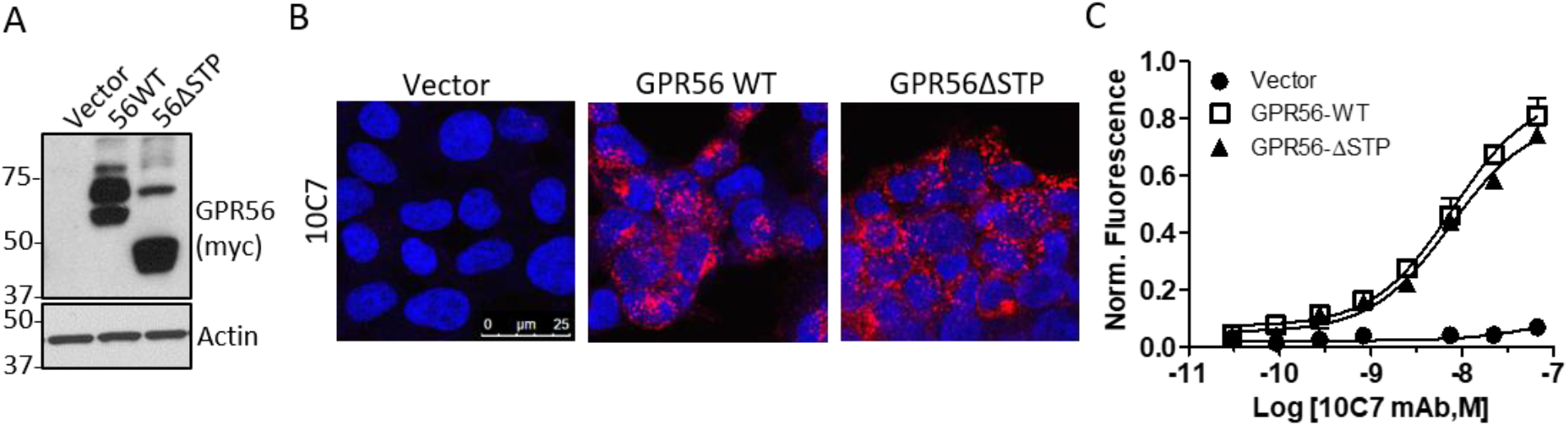
Novel Anti-GPR56 mAb 10C7 binds GPR56 with high specificity. A, Western blot of vector, GPR56WT, and mutant GPR56ΔSTP stable expression in 293T cells. B, Confocal microscopy images of 10C7 (5 μg/ml) binding and internalization in stable cell lines after 1hr treatment at 37°C C, Cell-based binding assay shows 10C7 binds WT and ΔSTP, but not vector cells in a concentration-dependent manner. Error bars are S.E. (n=3).

### GPR56 overexpression activates SRF-RE independent of the STP region of the ECD

GPR56 has been shown to induce activation of the small GTPase RhoA mediated through Gα_12/13_ to promote cell adhesion (13,20,24–26). To examine if mutant GPR56 ΔSTP was fully functional, we utilized a GTPase pulldown assay that employs the Rho binding domain of the Rho effector, Rhotekin. Western blot analysis showed that overexpression of GPR56 WT or GPR56 ΔSTP in 293T cells equivalently increased active RhoA (RhoA-GTP) levels compared to vector control (Fig. 2A). To measure downstream signaling effects, we employed the SRF-RE reporter which drives transcription of a luciferase reporter gene in response to G_12/13_-RhoA-mediated activation of SRF signaling. SRF binding to its response element promotes transcription of target genes involved in the regulation of cell adhesion and actin dynamics (28). A series of reports have shown that GPR56 can activate both SRF-RE (20) and the SRE reporter (21,23,29). SRF-RE is similar to SRE, yet is a more precise readout for G_12/13_-RhoA-mediated signaling (30). Consistent with active RhoA levels, both GPR56 WT and ΔSTP significantly increased SRF-RE activity (approximately 6-fold) compared to vector (Fig. 2B). Treatment with a Rho inhibitor suppressed both SRF-RE response and active RhoA levels (Figs. 2B and S1C), confirming that GPR56-mediated SRF-RE activity is mediated through RhoA. These data show that the STP region is not essential for constitutive GPR56-mediated RhoA signaling.

**Figure 2.**
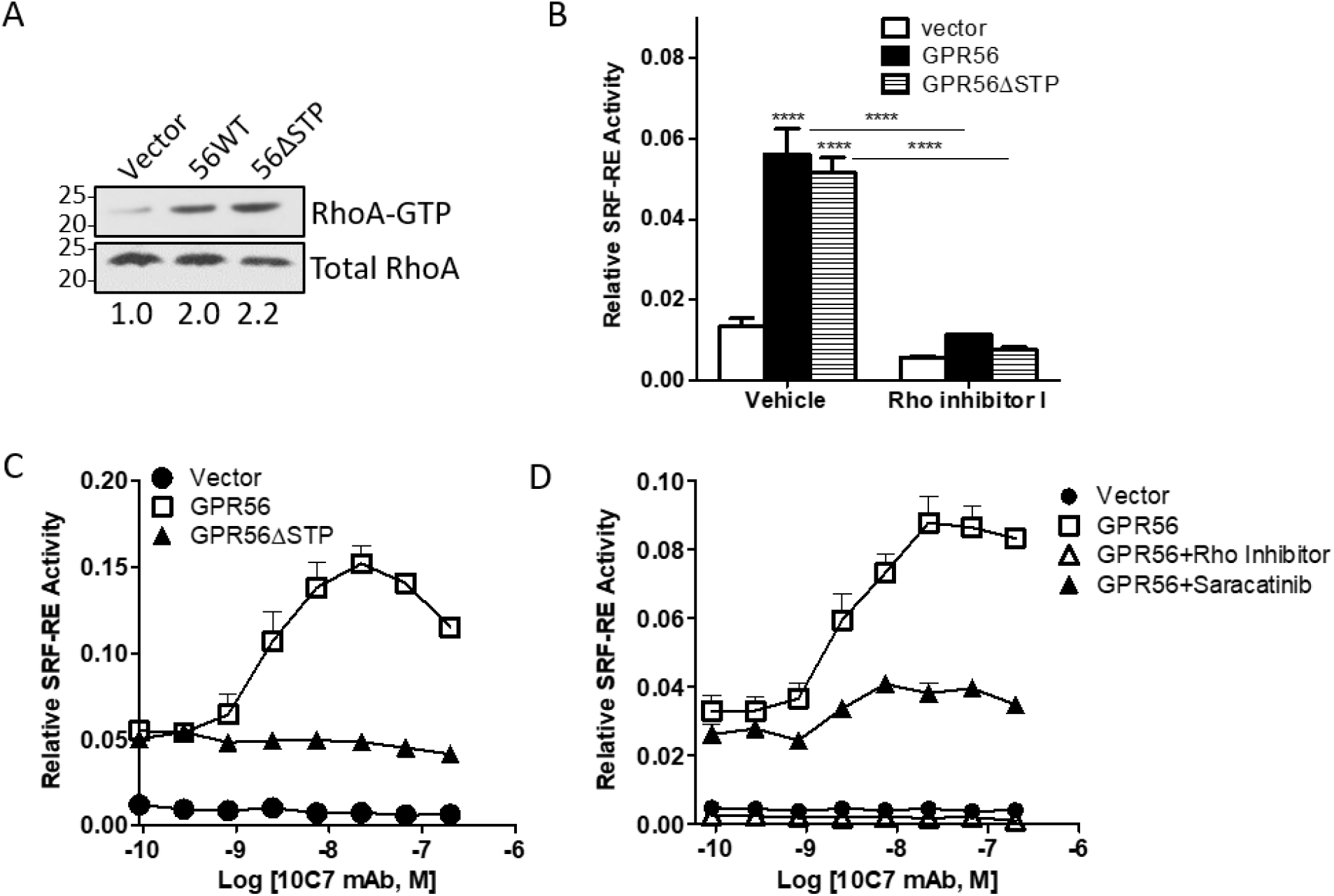
10C7 potentiates GPR56 activation of RhoA-mediated SRF signaling. A, RhoA activation assay shows increased levels in GPR56WT and ΔSTP cells compared to vector cells. B, Overexpression of GPR56 WT and ΔSTP enhances SRF-RE activity, which is blocked by pre-treatment with Rho inhibitor I (2μg/ml) for 3hrs. Statistical significance determined by ANOVA, **** p<0.0001. C, 10C7 induces a dose-dependent potentiation of SRF-RE in GPR56WT, but not ΔSTP cells. D, Effects of Rho inhibitor I (2ug/ml) and Src inhibitor (saracatinib) on 10C7 potentiation of GPR56-meditated SRF signaling. Error bars are S.E. (n=3).

### 10C7-induced SRF-RE activity is dependent on the STP region and activation of Src

Previously we and others have demonstrated that mAbs targeting 7TM receptors can function to activate intracellular signaling (31–33). Therefore, we tested whether 10C7 could modulate GPR56-mediated SRF-RE activity. 293T cells were co-transfected with the vector control or GPR56 expression plasmids and SRF-RE then treated with 10C7 in a dose-dependent manner. Interestingly, we found that 10C7 induced SRF-RE activity by approximately 3-fold in GPR56WT with an EC50 of ~2nM, but not in ΔSTP cells (Fig. 2C). No 10C7-induced effects were observed in vector transfected cells. These findings suggest that 10C7 could act as an agonist on GPR56 to trigger SRF-RE signaling in a STP-dependent fashion.

To identify potential new molecular players contributing to 10C7-induced potentiation of GPR56-mediated RhoA-SRF activity, we screened a variety of chemical inhibitors utilizing the SRF-RE assay. As expected, treatment Rho inhibitor completely suppressed constitutive and 10C7-induced SRF-RE response in GPR56 WT cells (Figs. 2D) and basal response in vector cells (Fig. S1D). Interestingly, we found that treatment with saracatinib, an inhibitor of the Src family of tyrosine kinases (Src), suppressed 10C7 potentiation of the SRF-RE reporter, yet only had minimal effect on baseline activity in GPR56WT cells (Fig. 2D). Saracatinib has no effect on SRF-RE activity in vector cells (Fig. S1D). These results demonstrate that Src activation may play a role in 10C7-induced activation of GPR56-mediated SRF signaling.

### GPR56 activates Src-Fak adhesion signaling and is potentiated by 10C7

Since Src and focal adhesion kinase (FAK) are widely associated with adhesion signaling, we tested if overexpression of GPR56 could modulate Src-Fak activity. As shown in Fig. 3A, GPR56WT stable cells showed increased phosphorylated levels of Src, Fak and paxillin (Pax) compared to vector. This was also confirmed by transient transfection (Fig. S2A). Furthermore, overexpression of GPR56ΔSTP had no effect on Src-Fak phosphorylation. Treatment of cells with 3μg/ml (~20 nM) 10C7 considerably enhanced phosphorylation of Src, Fak, and Pax in GPR56WT cells but not in ΔSTP cells (Fig. 3A). Dose-dependent treatment of GPR56WT cells showed maximum phosphorylation was induced by 3 μg/ml 10C7 after 1 hour. No change in phosphorylation was detected after treatment of GPR56 WT cells with a non-targeting isotype control antibody (hIgG1) or an anti-myc-tag mAb which binds to the N-terminus of recombinant GPR56WT (Fig. 3C and S2B). Time-dependent treatment of GPR56WT cells with 3μg/ml showed increased phosphorylation of Src, Fak, and Pax at 5 minutes with peak phosphorylation of Src observed at 15 minutes post-treatment (Fig. 3D). Elevated phosphorylation levels were still detected after 6 hours post-treatment with 10C7 and no significant changes in phosphorylation was detected in vector cells (Fig. S2C). Since GPR56 drives cell adhesion, we investigated if pre-treatment with 10C7 could enhance adhesion to a collagen in a GPR56-specific manner. Our results show that GPR56 expression increased adhesion of 293T cells compared to vector, with approximate 2-fold increase observed after 10 min (Fig. 3E). Furthermore, 10C7 treatment significantly increased the rate of adhesion of GPR56WT cells after 15 min, yet had no effect on vector cells. These findings demonstrate that 10C7 can potentiate GPR56-mediated Src-Fak signaling and cell adhesion in 293T cells.

**Figure 3.**
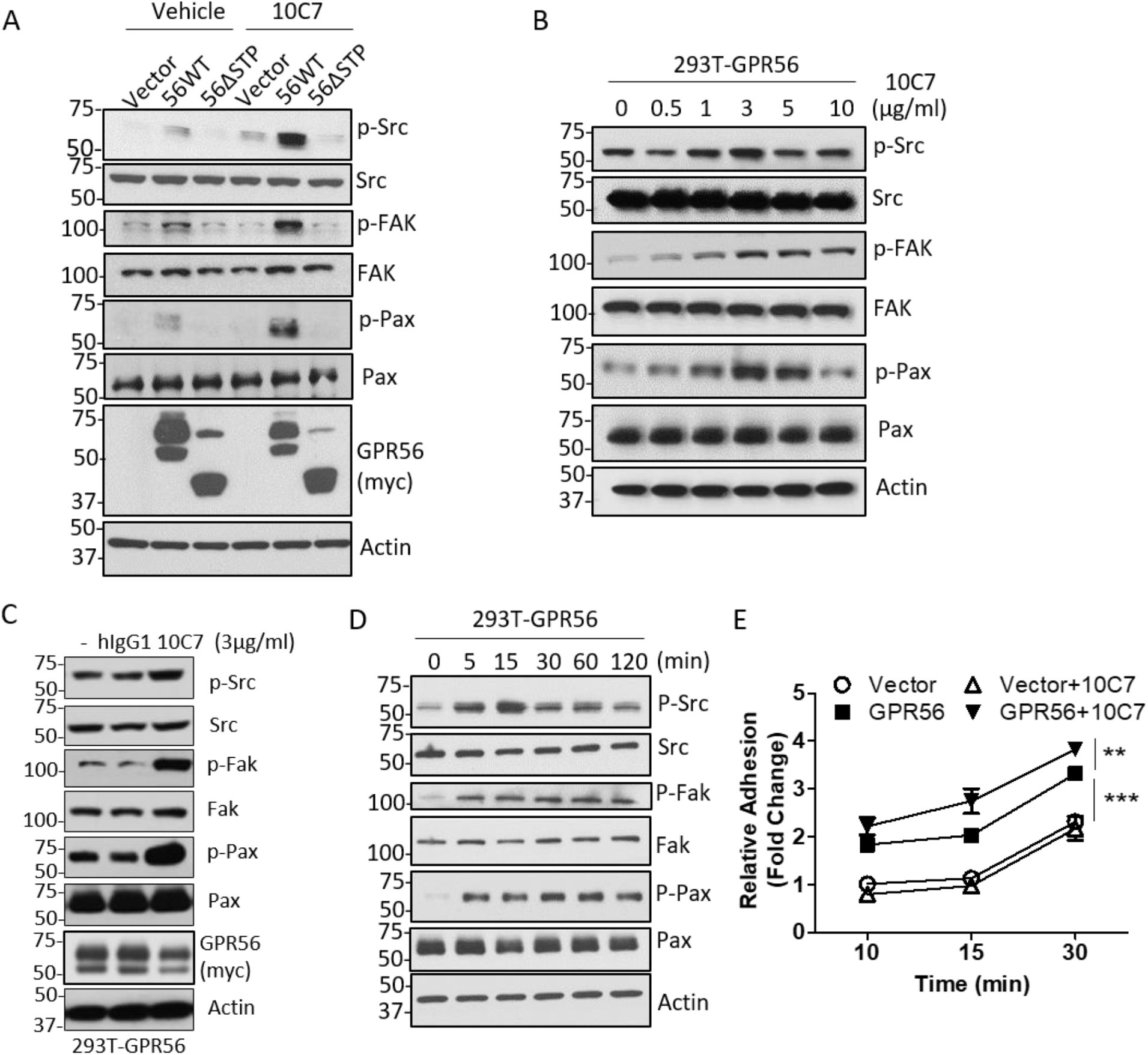
GPR56 activates Src-Fak adhesion signaling in 293T cells and requires the STP domain. A, Western blot showing GPR56WT, but not ΔSTP, activates phosphorylation of Src, Fak, and paxillin which is further enhanced by treatment with 10C7 (3μg/ml) for 1 hr. B, 10C7 dose-dependent phosphorylation in serum starved 293T-GPR56 cells treated for 1 hr. C, 10C7 and not isotype control antibody (hIgG1) potentiates Src-Fak signaling in GPR56 transfected cells after 30 min. D, Time-dependent effects on phosphorylation in serum starved 293T-GPR56 cells treated with 10C7 (3μg/ml). E, 293T-GPR56 cells exhibit increased adhesion to collagen, which is augmented by 10C7. Statistical significance determined by 2-way ANOVA, ** p<0.01, and ***p<0.001. All error bars are S.E. (n=3).

### Depletion of GPR56 decreases Src-Fak phosphorylation and adhesion of colorectal cancer cells

Similar to GPR56, Src and Fak are upregulated in CRC and high expression correlates with poor clinical prognosis (12,13). Therefore, we investigated if 10C7 could specifically bind endogenous GPR56 to activate Src-Fak signaling in CRC cell lines. Previously, we reported that DLD-1 and HT-29 cells express high levels of GPR56 and we established GPR56 shRNA knockdown (KD) cell lines (13). ICC and confocal analysis showed 10C7 binding and internalization in DLD-1 cells after 1 hr at 37°C (Fig. 4A). No binding of the hIgG1 isotype control was detected. Additionally, 10C7 did not bind DLD-1 GPR56 KD (shGPR56) cells, yet shRNA control (shCTL) cells showed similar binding and internalization as observed in parental DLD-1 cells (Fig. 4B). These results validate 10C7 binding specificity for endogenous GPR56. We then tested if treatment of CRC cells with 10C7 could activate Src-Fak signaling. Cells were serum starved and treated with 25 μg/ml 10C7 at the indicated time-points. Interestingly, both cell lines showed increased levels of Src, Fak, and Pax phosphorylation at 30 min that returned to baseline levels or below at 2 hrs (Fig 4C). To determine if ablation/depletion of endogenous GPR56 expression affects Src-Fak signaling, we performed western blot using our previously established shCTL and shGPR56 DLD-1 and HT-29 cell lines(13). GPR56 KD using two different shRNAs reduced phosphorylated levels of Src, Fak, and Pax, indicating that endogenous GPR56 mediates Src-Fak signaling exclusive of 10C7 treatment (Fig. 4D). We then tested if GPR56 KD effected cell adhesion. As shown in Figs. 4E-F, loss of GPR56 expression in both DLD-1 and HT-29 cells significantly decreased the rate of cell adhesion to a collagen matrix. These data demonstrate that GPR56 can regulate Src-Fak adhesion signaling in CRC cells.

**Figure 4.**
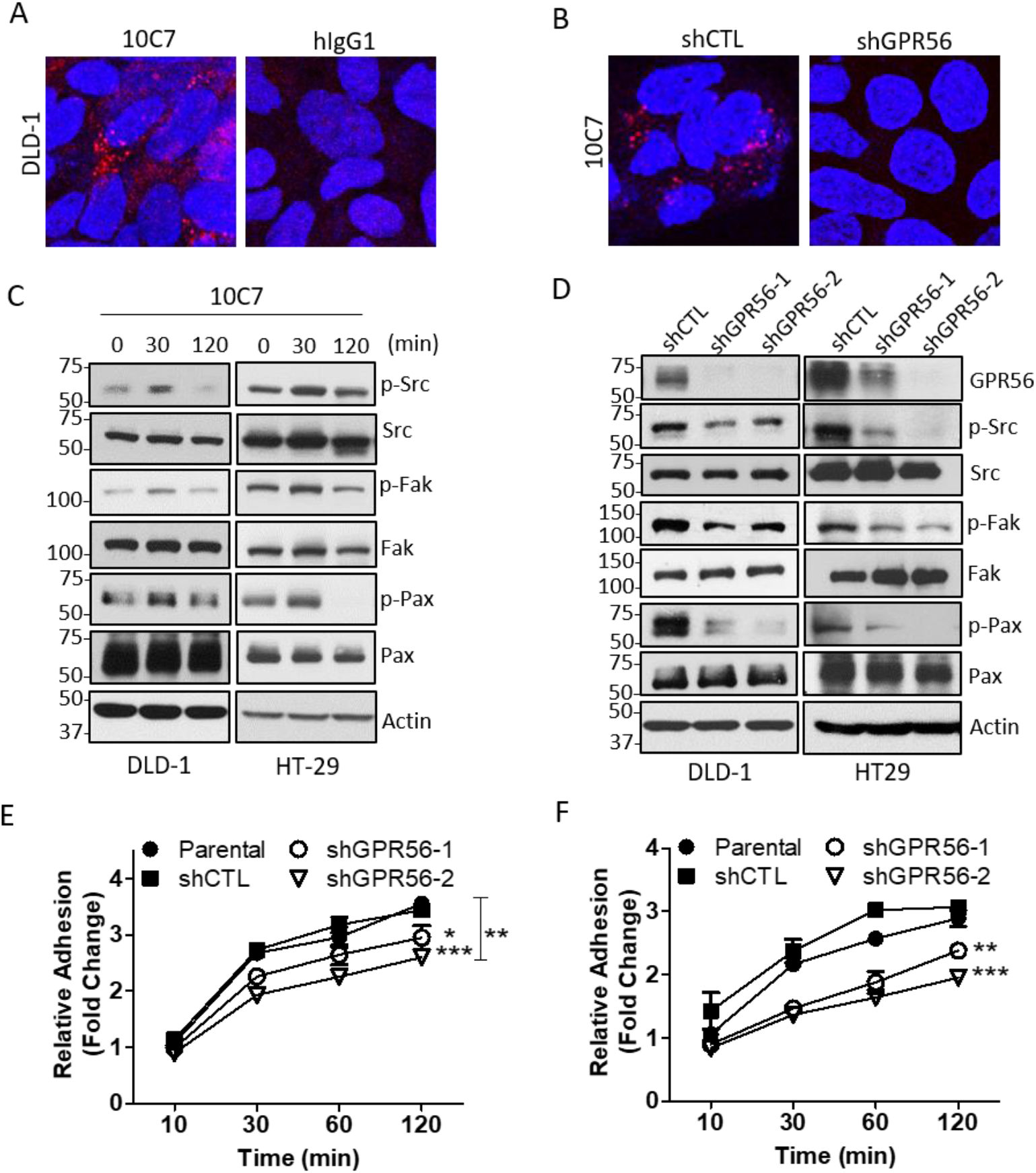
GPR56 regulates Src-Fak phosphorylation and adhesion in colorectal cancer cells. A, Confocal microscopy images of ICC showing 10C7 (15 μg/ml) binds GPR56 and internalizes in DLD-1 cells after 1 hr at 37°C. No binding was detected using non-targeting hIgG1 isotype control. B, 10C7 binds shRNA control (shCTL), but not GPR56 shRNA knockdown DLD-1 cells (shGPR56-2). C, Western blot showing time-dependent effects of 10C7 (20 μg/ml) treatment on phosphorylation of Src, Fak, and paxillin in serum starved DLD-1 and HT-29 cells. D, GPR56 knockdown decreases phosphorylation of Src, Fak, paxillin and E-F, reduces adhesion to collagen in E, DLD-1 and F, HT-29 cells. Statistical significance determined by 2-way ANOVA. For DLD-1 cells * p<0.05, and ***p<0.001 vs. shCTL and ** p<0.01 for shGPR56-2 vs. parental cells. For HT-29 cells, ** p<0.01, and ***p<0.001 vs. shCTL and parental cells. Error bars are S.E. (n=3).

### GPR56-mediated Src-Fak phosphorylation is independent of RhoA activation

Since we found that 10C7 potentiation of GPR56-mediated SRF-RE signaling required Src, we wanted to determine if phosphorylation of Src, Fak, and Pax was dependent on Gα_12/13_ and activation of RhoA. 293T cells were co-transfected with GPR56 and either Gα_13_-targeting or control siRNA and treated with or without 10C7. As shown in Fig. 5A, knockdown of Gα_13_ did not have a significant effect on baseline or 10C7-induced phosphorylation mediated by GPR56. We then examined if RhoA activation is required for GPR56-mediated phosphorylation of Src, Fak, and Pax. 293T-GPR56 cells were pre-treated with Rho inhibitor for 3 hrs and then treated with or without 10C7. Results showed that treatment with Rho inhibitor failed to suppress phosphorylation (Fig. 5B), suggesting that GPR56-mediated activation of RhoA is not essential for activation of Src-Fak signaling. In fact, pre-treatment with the Rho inhibitor lead to a slight elevation in baseline p-Src levels. Next, we investigated whether Src or Fak phosphorylation occurs immediately following GPR56 activation. We pre-treated 293T-GPR56 cells with either Src inhibitor (saracatinib) or Fak inhibitor (defactinib) for 3 hrs, then treated with or without 10C7 (Figs. 4B-C). As shown in Fig. 5B, saracatinib inhibited both baseline and 10C7-induced phosphorylation of Src, Fak, and Pax. Treatment with Src inhibitor, PP2, showed similar effects (Fig. S3). Defactinib treatment lead to inhibition of p-Fak, as expected, and 10C7-induced p-Pax (Fig. 5C). No inhibition of Src phosphorylation was observed and only a partial inhibition of baseline p-Pax levels was detected (Fig. 5C). These data indicate that GPR56-mediated Src phosphorylation occurs upstream of Fak phosphorylation and that both Src and Fak can potentially phosphorylate Pax. Moreover, our findings suggest that GPR56-mediated Src-Fak signaling does not require activation of Gα_13_-RhoA.

**Figure 5.**
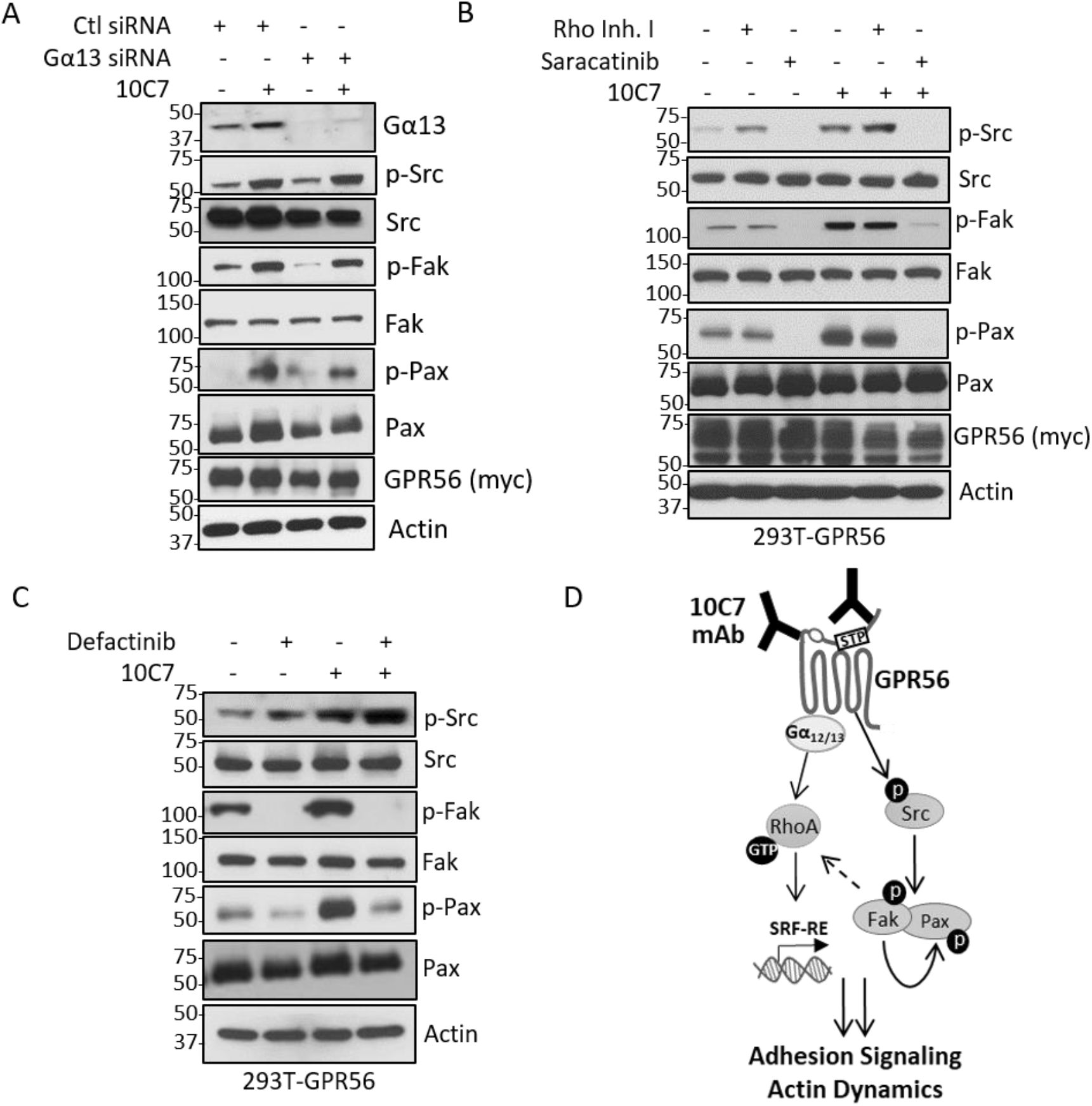
GPR56-mediated phosphorylation of Src occurs upstream of Fak and is independent of RhoA activation. A, Western blot showing Gα_13_ knockdown does not significantly effect GPR56-mediated Src/Fak signaling in 293T cells. (B-C), Western blots showing effect of inhibitors of B, RhoA (Rho Inhibitor I, 2μg/ml), Src (saracatinib, 10μM), and C, Fak (defactinib, 10μM) on GPR56 constitutive and 10C7-induced phosphorylation. Stable 293T-GPR56 WT cells were pre-treated with inhibitors for 3 hrs, then treated with 10C7 (3μg/ml) for 45 min. D, Schematic of 10C7 mAb-induced GPR56 adhesion signaling model. GPR56 activates Gα_3_-RhoA-SRF signaling and promotes Src phosphorylation independent of RhoA. Src activation leads to phosphorylation of Fak and paxilllin. Binding of 10C7, or possibly a natural ligand, to an unidentified position on the ECD induces a conformational change of the ECD to potentiate Src-Fak signaling. The STP site of the ECD is not required for 10C7 binding, but is essential for activation of Src-Fak. Potentiation of Src-Fak signaling enhances RhoA-SRF signaling downstream of G_13_. GPR56 likely coordinates activation of these signaling pathways to regulate adhesion and other cellular processes.

## Discussion

GPR56 has been shown to have essential functions in many physiological processes and emerging evidence has established a critical role for GPR56 in tumor progression (2–4,10,12–15). However, due to its intricate structure and lack of high affinity receptor-specific ligand(s), the signaling mechanism of GPR56 still remains poorly understood. In this study, we generated a unique anti-GPR56 mAb, 10C7, directed against the GPR56 ECD in order to interrogate its signaling mechanism(s), particularly with endogenously expressed receptor. Based on our findings, we propose a model by which GPR56 can activate both G_12/13_-RhoA-SRF and Src-Fak signaling to regulate adhesion in normal and cancer cells (Fig. 5D). Treatment with 10C7, or potentially an endogenous ligand, can potentiate Src phosphorylation leading to subsequent phosphorylation of Fak and Pax. GPR56-mediated Src-Fak signaling requires the STP region of the NTF which may be involved in a conformational shift of the receptor. This conformational change may promote association of the NTF with the stalk region, an extracellular loop of the TM, and/or potentially a co-receptor to transmit signaling. Interestingly, Jeong et al demonstrated synergistic activities of GPR56 and α3β1 integrin during cerebral cortical development (34), which may account for GPR56 activation of integrin partners such as Src-Fak. Additionally, GPR56-interacting proteins including CD81 and collagen III (24,35) have also been shown to associate with integrins (36–38). The STP region overlaps with the PLL domain which shares sequence similarity to LNS domains which can mediate cell adhesion (19,39). Deletion of residues within this region have been shown to inhibit GPR56 interaction with extracellular matrix proteins (7,24). This is consistent with our finding that GPR56ΔSTP fails to activate Src-Fak phosphorylation and suggests that this region plays an important role in regulating adhesion signaling. Furthermore, we showed that 10C7-induced Src-Fak signaling enhanced RhoA-SRF signaling through an unknown mechanism. Previously, Iguchi et al also showed that a polyclonal anti-GPR56 antibody could induce RhoA activation and SRE-mediated reporter activity (25). Several studies have demonstrated that Rho guanine nucleotide exchange factors (RhoGEFs) can bind and function as substrates for Fak (40–42). Fak activation of RhoGEFs could play a role in controlling active RhoA levels downstream of Src, thus leading to increased SRF signaling. The manner by which these GPR56-mediated signaling pathways function to coordinate the regulation of actin dynamics and cell adhesion requires further investigation.

Different stalk-dependent and -independent models have been proposed to provide structural and mechanistic insight into how extracellular interactions with GPR56 trigger intracellular signaling activity. However, it is possible that these models function in concert with each other or the GPR56 activation mechanism(s) may essentially be ligand- and/or pathway-dependent. We showed that 10C7 co-internalized with GPR56, which initially suggested two potential scenarios of receptor activation; (1) 10C7 binds the NTF of the ECD and NTF dissociation is not required for potentiation of Src-Fak signaling or (2) 10C7 directly binds the stalk region of the CTF and NTF dissociation may or may not occur (Fig. 1C). A similar internalization pattern was shown using a commercial mAb which binds the myc-tag at the N-terminal of recombinant GPR56, though this mAb did not activate Src-Fak phosphorylation (Figs. S1B and S2B). However, we showed that the STP region of the NTF was not required for 10C7 binding (Fig. 1B-C). These data indicate that 10C7 directly binds to either the N-terminus of the PLL domain, the GAIN domain, or the stalk region, yet this interaction is not sufficient for receptor activation of Src-Fak signaling. We theorize that a 10C7-induced conformational change which promotes association of the ECD, specifically the STP region, with the CTF and/or co-receptor may be obligate to activate Src-Fak signaling. Thus, GPR56 activation of Src-Fak signaling appears to be stalk-independent. Consistently, Ohta et al generated GPR56 mAbs and showed that agonistic mAbs could enhance interaction of NTF with the CTF (43). However, they utilized activation of G_q_ and inhibition of cell migration as a readout for agonist activity. Furthermore, the use of monobodies targeting different regions of the ECD supports stalk-independent regulation of GPR56-mediated SRE activity (23). Based on findings by Kishore et al, GAIN domain cleavage of the NTF was not necessary for GPR56-mediated basal activation of the SRF reporter, yet removal of the CTF stalk region significantly abrogated SRF reporter activity (20). Thus, the role that the residues of stalk region play in GPR56-mediated Src-Fak signaling and potentiation of RhoA-SRF remains to be determined. Comprehensive epitope mapping of 10C7 will be required to further elucidate this mechanism.

Expression levels of Src and Fak are elevated in CRC and increased Src-Fak activation has been implicated in tumor growth and metastasis (44–48). Similarly, GPR56 is highly upregulated in CRC and shown to promote proliferation of CRC cells and tumor growth in vivo. Therefore, we utilized CRC cell lines to investigate endogenous effects of GPR56 on Src-Fak signaling. Similar to 293T cells, 10C7 potentiated Src-Fak phosphorylation in CRC cell lines and GPR56 knockdown suppressed Src-Fak signaling and cell adhesion (Figs. 4C-F). Previous studies have demonstrated that loss of GPR56 in CRC cells lead to decreased migration, invasion, and resistance to chemotherapy (12,13,16); all of which may potentially be regulated through the Src-Fak pathway. In fact, therapeutic inhibitors of Src have been shown to suppress CRC cell growth and adhesion and sensitize cells to chemotherapy (49–51). Moreover, both high GPR56 expression and activated Src-Fak have been associated with poor prognosis in CRC (12,13,44,52). In melanoma, where GPR56 plays an inhibitory role, Millar et al showed by ICC that GPR56 depletion lead to increased phosphorylation of Fak (53). This suggests that GPR56 may have opposing roles in different cancers and may be dependent on differential expression of its natural ligands or crosstalk with other signaling pathways. Based on these findings, it will be of interest to further explore the role of GPR56-mediated Src-Fak signaling in the regulation of tumor progression and drug resistance.

In conclusion, we generated a high affinity anti-GPR56 mAb that can function as a biological tool to delineate the novel activation mechanism and signaling pathways of GPR56. Our findings uncover a unique ECD-dependent mechanism by which GPR56 potentially activates Src-Fak signaling to regulate normal and pathological processes. Therapeutics targeting GPR56 to suppress Src-Fak signaling could hold significant potential for the treatment of cancers with high expression of GPR56, particularly CRC.

## Experimental Procedures

### Plasmids and cloning

Reporter vector pGL4.34[luc2P/SRF-RE/Hygro] was purchased from Promega. The myc-tagged GPR56 wild-type (GPR56 WT) vector (pIRESpuro3-myc-hGPR56) encoding amino acids 26-693 was previously described (13) and the mutant GPR56ΔSTP vector (pIRESpuro3-myc-hGPR56ΔSTP) lacking the STP domain was generated in a similar manner. Briefly, the sequence encoding GPR56ΔSTP (amino acids 26-693 with deletion of 108-177 within the ECD) was subcloned from BC-deltaSTP-GPR56 and fused with sequences encoding a Myc tag at the N terminus, and cloned downstream of a sequence encoding the CD8 signal peptide (MALPVTALLLPLALLLHAA) in the vector pIRESpuro3 (Clontech). BC-deltaSTP-GPR56 was from Lei Xu (Addgene, 44198). For antibody generation, the sequence encoding the native signal peptide and ECD of GPR56 (amino acids 1-400) was subcloned from pCAG-hGPR56-IRES-GFP (from Christopher A Walsh, Addgene 52297) into pCEP4 (Invitrogen) and fused to a C-terminal 6x His-tag (GPR56ECD-6xHis).

### Commercial antibodies, chemical inhibitors, and other reagents

Commercial antibodies were used in accordance to manufacturer’s guidelines. For western blot: GPR56 (H00009289-B01P) from Abnova, pSrcY416 (6943), Src (2123), pFakY397 (8556), Fak (3285), p-paxillinY118 (2541), β-actin (3700), myc-tag (2276) from Cell Signaling, and paxillin (610863) from BD Biosciences. For immunocytochemistry (ICC) and cell-based binding assays: secondary goat anti-human-Alexa-555 (Life Technologies). The cell-permeable C3 transferase-based Rho inhibitor 1 was from Cytoskeleton. Saracatinib and defactinib were from Selleck Chemicals. hIgG1 isotype control was from Fisher Scientific.

### Cell culture, transfection, and stable cell line generation

HEK293T (293T), DLD-1, and HT-29 cells were purchased from ATCC. Cell lines were authenticated utilizing short tandem repeat profiling, routinely tested for mycoplasma. 293T cells were cultured in DMEM and colon cancer cell lines in RPMI medium and supplemented with 10% fetal bovine serum and penicillin/streptomycin. Transient transfections were performed in 6-well plates (1.5μg DNA/well) using jetPRIME (Polypus Transfection). Control and Gα_13_-targeting SMARTtpool siRNA M-009948-00-0005 (Horizon Discovery) was transfected at a final concentration of 50nM. To generate stable 293T cell lines, cells were transfected with GPR56 WT, GPR56ΔSTP, or control vector and selected in 1 μg/ml puromycin. DLD-1 and HT-29 shRNA control (pLKO.1, shCTL) and GPR56 (shGPR56) knockdown cell lines were generated by lentiviral infection as previously reported (13).

### Generation of anti-GPR56 monoclonal antibody

The anti-GPR56 mAb, 10C7, was generated together with ProMab Biotechnologies (Richmond, CA, USA, promab.com). Briefly, GPR56ECD-6xHis was purified using a Ni-NTA column and mice were immunized with the purified ECD. After a series of immunization and cloning steps, hybridoma clones were selected and the supernatants screened for binding specificity by ELISA and immunocytochemistry (ICC). The mouse variable heavy and light chain (VH and VL) sequences were amplified by PCR, cloned, and then sequenced. To produce a mouse-human chimera 10C7 mAb, the mouse VH and VL were subcloned into pCEP4 expression vectors containing the constant region of human IgG1 heavy chain (CH) and kappa light chain (CL) as previously described (54). Large-scale mAb production and purification was performed as reported earlier (31).

### Western blot and RhoA pulldown assays

RhoA-GTP pulldown activation assays (Cytoskeleton, #BK036) were carried out according to protocol. For western blots, protein extraction was performed using RIPA buffer (Sigma) supplemented with protease/phosphatase inhibitors. Cell lysates were incubated at 37°C for 1 hr and diluted in laemmli sample buffer prior to loading on SDS-PAGE. HRP-labeled secondary antibodies were utilized for detection with the standard ECL protocol. Quantification of RhoA was performed using ImageJ.

### Immunocytochemistry and cell-based binding assays

For immunocytochemistry (ICC), HEK293T or DLD-1 cells were seeded into 8-well chamber slides and incubated overnight. The next day, cells were treated with 10C7 or hIgG1 isotype control at 37°C for 1 hr. Cells were washed with PBS, fixed in 4% formalin, permeabilized in 0.1% saponin, then incubated with goat anti-human-Alexa-555 for 1 hr at room temperature. Nuclei were counterstained with TO-PRO^®^-3. Images were acquired using confocal Leica TCS SP5 microscope with LAS AF Lite software. Whole cell-based binding assays were performed as previously described (54). Fluorescence intensity was quantified using Tecan Infinite M1000 plate reader.

### Luciferase reporter assays

Cells were plated at 2500 cells/well in 96 half-well plates. Serial dilutions of 10C7 and/or chemical inhibitors were added and allowed to incubate at 37°C overnight. Luciferase activity was measured using Pierce™ Firefly Luciferase Glow Assay Kit according to manufacturer’s protocol using an EnVision mulitlabel plate reader (PerkinElmer). AlamarBlue (ThermoFisher) was used for normalization to cell viability and fluorescence quantified at 530/590 nm using Tecan Infinite M1000 plate reader. Each condition was performed in triplicate, n ≥ 3.

### Adhesion assays

Cell were seeded in a 96-well collagen I-coated plate at 15,000 cells/well (1.5×10^5^ cells/ml), in the appropriate culture medium and incubated at 37°C for the indicated time-points. Non-attached cells were washed with PBS, fixed with 4% formalin for 15 min and labeled with 5 mg/ml crystal violet for 10 min. Plates were washed with PBS to remove unbound dye. Crystal violet from adherent cells was solubilized using 2% SDS and absorbance was measured at 595 nm using Tecan Infinite M1000 plate reader.

### Statistical analysis

Statistical analysis was performed using GraphPad Prism software. Data are expressed as mean +/- standard error of the mean (S.E.) and for all experiments n ≥ 3. K_d_ values were determined using logistic nonlinear regression model. For adhesion assays, differences between groups were analyzed by two-way ANOVA. Other multiple comparisons used one-way ANOVA and Tukey’s post-hoc analysis. *P* ≤ 0.05 was considered statistically significant.

### Data availability

All data are contained within the article.

## Funding and additional information

This work was supported by funding from the National Institutes of Health (R01CA226894) and Welch Foundation Endowment Fund Award (L-AU-0002-19940421) to K.S. and National Institutes of Health (R01GM102485) and Cancer Prevention and Research Institute of Texas (RP170245) to Q. J. Liu.

## Conflict of Interest

The authors declare no potential conflicts of interest.

## Abbreviations

The abbreviations used are:

GPR56: G protein-coupled receptor 56
ADGRG1: adhesion G protein-coupled receptor G1
aGPCR: adhesion G protein-coupled receptor
mAb: monoclonal antibody
ECD: extracellular domain
7TM: seven-transmembrane
WT: wild-type
NTF: N-terminal fragment
CTF: C-terminal fragment
PLL: Pentraxin/Laminin/neurexin/sex-hormone(LNS)-binding-globulin-Like
STP: SerineThreonine-Proline-rich
GAIN: GPCR-autoproteolysis inducing
GPS: GPCR Proteolysis Site
Fak: focal adhesion kinase
Src: Src proto-oncogene, non-receptor tyrosine kinase
Pax: paxillin
KD: knockdown
shRNA: short hairpin ribonucleic acid
siRNA: small interfering ribonucleic acid
ICC: immunocytochemistry
SRE: serum response element
SRF-RE: serum response factor response element
RhoA: Ras homolog family member A
CRC: colorectal cancer

